# How to optimally sample a sequence for rapid analysis

**DOI:** 10.1101/2022.08.18.504476

**Authors:** Martin C. Frith, Jim Shaw, John L. Spouge

## Abstract

We face an increasing flood of genetic sequence data, from diverse sources, requiring rapid computational analysis. Rapid analysis can be achieved by sampling a subset of positions in each sequence. Previous sequence-sampling methods, such as minimizers, syncmers, and minimally-overlapping words, were developed by heuristic intuition, and are not optimal.

We present a sequence-sampling approach that provably optimizes sensitivity for a whole class of sequence comparison methods, for randomly-evolving sequences. It it likely near-optimal for a wide range of alignment-based and alignment-free analyses. For real biological DNA, it increases specificity by avoiding simple repeats. Our approach generalizes universal hitting sets (which guarantee to sample a sequence at least once), and polar sets (which guarantee to sample a sequence at most once). This helps us understand how to do rapid sequence analysis as accurately as possible.

## Introduction

Let us start with a curious puzzle. Suppose we randomly pick one of the 4^5^ DNA sequences of length 5. Is the probability that it contains the substring `acgt` equal to the probability that it contains `aaaa`?

There seems to be a strong intuition that the answer is “yes”: let us check. There are four length-5 sequences that start with `acgt`, and another four that end with `acgt`, for a total of eight. On the other hand, there are seven sequences containing `aaaa`: `aaaaa` plus three others starting with `aaaa` and three others ending with `aaaa`. It is remarkable that such a simple puzzle can be so counterintuitive.

This curious property of sequences is not just a shallow trick, because it can be leveraged into useful methods. For example, it is the basis of spaced seeds [17]. Here, we build upon it a theory of optimal sequence-sampling, for rapid analysis of big data.

The main way of analyzing genetic sequences is by comparing them to each other. For large data, this is usually done via “seeds”, by which we mean simple similarities that can be found quickly [31]. The simplest seeds are fixed-length exact matches, but they can also be inexact [1, 17, 22, 28] and/or variable-length [8]. These seeds are used in diverse ways to infer relationships between sequences.

The seed-and-extend approach extends seeds into detailed alignments. The simplest “one hit” method attempts extension from each seed [1]. A “two hit” alternative triggers extension from a pair of nearby, non-overlapping seeds [2]. Another “coverage” method triggers extension from a group of possibly-overlapping seeds, if the number of letters covered by the seeds exceeds a threshold [22, 3, 23, 21].

There are also alignment-free approaches [32], which often use seeds, also called “micro-alignments” [34]. These methods are used to infer evolutionary relationships, e.g. by estimating evolutionary distances between sequences [5, 19]. For example, they are used to classify sequences taxonomically, based on seed matches to a database of reference genomes [26, 33], which is phylogenetic placement [15].

For faster or larger-scale sequence comparison, perhaps the most promising approach is sampling, for example just use seeds starting with `a`. Sampling methods include: anchor strings [18], minimizers [29, 27], syncmers [10], and minimally-overlapping words [12]. Those studies showed that some sampling methods are better than others in some situations.

Previous sampling methods, however, were developed by heuristic intuition, and are unlikely to be optimal. It is also possible that a sampling method is superior in some scenarios but not others, e.g. for closely-but not distantly-related sequences, or for alignment-based but not alignment-free methods. Thus, we lack a non-heuristic criterion to optimize sampling.

Here we develop sampling methods, for exact-match seeds, which provably optimize sensitivity for one-hit-based sequence comparison, for randomly-evolving sequences. We argue that they are likely near-optimal in a wide range of alignment-based and alignment-free scenarios. We then show evidence that they work well for real biological DNA. The main advance over previous heuristics [12] is improved theoretical understanding of how to sample positions in a sequence near-optimally for a wide range of applications.

### Illustration of our sampling approach

The main idea is that related sequences have runs of overlapping matches (fig. 1), and we maximize the probability of sampling these runs. To make things simple, we assume a random DNA sequence of independent, equally-probable bases, evolving by random substitutions. Let us consider two sampling schemes: matches starting with `ry`, and matches starting with `rr`. (`r` means puRine: `a` or `g`, and `y` means pYrimidine: `c` or `t`.) Each scheme samples 1 in 4 matches on average. A run of, say, 2 overlapping matches is hit when `ry` (or `rr`) occurs at the start of either match, i.e. when it occurs in a sequence of length 3. A quarter of all length-3 sequences start with `ry`, and a *different* quarter end with `ry` (because it’s impossible to start *and* end with `ry`), so the total hitting probability is 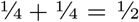. In contrast, 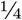 of length-3 sequences start with `rr` and an *overlapping* quarter end with `rr`, so the total hitting probability is lower (⅜). Thus, `ry` is more likely than `rr` to sample a run of 2 overlapping matches.

**Figure 1:**
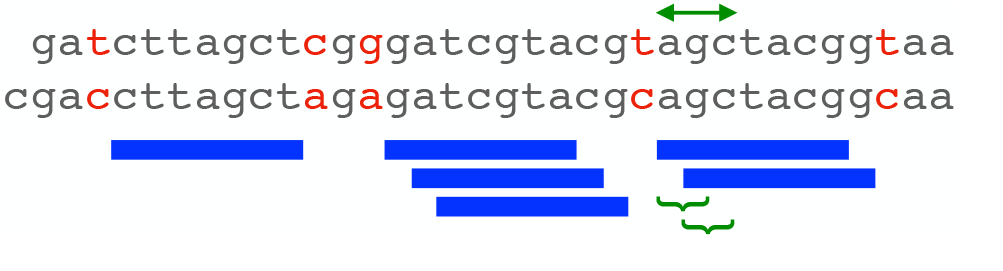
Example of related sequences, with length-7 exact matches (blue bars). The matches occur in runs of size 1, 3, and 2. If we sample matches starting with `ry`, we “hit” the size-2 run if `ry` is at either of the sites marked by braces, in other words, if `ry` occurs in the length-3 sequence marked by arrows.

In fact, `ry` is more likely than `rr` to sample a run of *x* overlapping matches, for any *x*≥ 2. This means that `ry` is more likely than `rr` to sample at least one match between two related sequences. Thus, `ry` is more sensitive, for one-hit-based sequence comparison. Moreover, it is more sensitive for any level of sequence divergence: for closely- or distantly-related sequences.

## Methods

### Sampling positions, matches, and runs

We sample positions in a sequence, and thereby sample matches between two sequences. For example, if we sample positions where `a` occurs, and then find matches starting at those positions, we will sample matches starting with `a`.

We are interested in sampling runs of overlapping matches (fig. 1). Sampling such a run is equivalent to sampling a run of consecutive starting positions of the matches in one sequence.

### Word-based sampling

A word-based sampling scheme specifies a set *Q* of length-*k* words, e.g. *Q* = {`rrry`, `ryrr`, `ryyr`, `yyyr}`, and selects the positions in a sequence where those words occur. This can be used to select seeds starting at those positions. We will focus on exact-match seeds of fixed length *m* ≥*k*. Some previous publications (e.g. [10]) have focused on the case *m* = *k*, so that both words and matches are *k*-mers, but here words are *k*-mers and matches are *m*-mers. Fundamentally, we are sampling positions in a sequence, and what we do with those positions is secondary.

Word-based sampling can be implemented with a lookup table of size |*A*| ^*k*^, where |*A*| is the alphabet size, to look up whether any word is in the set. This is fast and practical, as long as the table is small enough to fit in the computer’s fast cache memory. We use a purine/pyrimidine alphabet *A* = *{*`r`, `y`*}* because:

- In biological DNA, `r` and `y` have 50:50 frequencies, whereas `a`, `c`, `g`, and `t` have varying frequencies [6]. This makes the frequency of `r`/`y` words more predictable.
- Vertebrates have a high rate of `cg` →`tg`/`ca` substitutions [36]. This `cg` effect does not fit our model of independent letters with uniform substitution probability, but it does fit the model when the sites are encoded as `r`/`y` [4].
- A reduced alphabet makes it easier to use longer words, because the lookup table is smaller. Longer words can help to optimize sampling, by constraining the spacing between sampled positions.
- `r`/`y` word-based sampling is compatible with inexact seeds that allow `a` ↔`g` and `c`↔ `t` substitutions at some positions: such seeds often work well because these substitutions are frequent [22].

### Run-hitting probabilities

We wish to know the probability that a word-based sampling scheme hits a run of *x* consecutive start positions (fig. 1). This equals the probability that any word in our set occurs in a sequence of length *x* + *k* − 1.

This can be calculated by dynamic programming. Define *C*(*i, w*) to be the number of possible length-*i* sequences, with alphabet *A*, which have no substring that is in our set *Q*, and end in a length-*k* word *w*. The base case is:

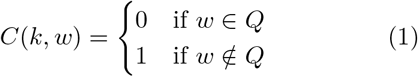

If we define *w*^*/*^ to be *w* with its final letter removed, and ⊕ is string concatenation:

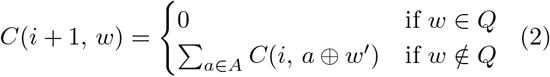

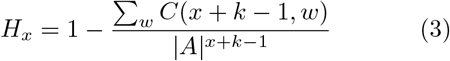

This dynamic programming method is similar to computation of hit probability for multiple spaced seed patterns [14].

### Upper bound

The run-hitting probabilities have a simple upper bound [30]. This is illustrated by the `ry` example in the Introduction. `ry` occurs with probability 1*/s* (*s* = 4 in this case), the probability of hitting a size-2 run is at most 2 *×* (1*/s*), and this upper bound is achieved when it’s impossible to sample both positions. In general, the probability of hitting a size-*x* run is at most: min(*x/s*, 1).

### Optimizing run-hitting probabilities

We wish to find a set of words with high run-hitting probabilities. We will try to maximize the *average* run-hitting probability, for runs of size 1, 2, 3, … *u*. When *u* is very large, this is equivalent to minimizing the variance of distance between consecutive sampled positions (see the Supplement).

We used two optimization methods: exact and heuristic. In both cases, we fix the word length *k* and the number of words *n*. The exact method evaluates all possible sets of *n* length-*k* words, and outputs all sets that have maximum average run-hitting probability. This is feasible only when *k* and *n* are small; we used it when *n* ≤8. The heuristic method uses simulated annealing (see the Supplement): it does not guarantee to find the maximum average run-hitting probability. These methods are available in maxhit-words from https://gitlab.com/mcfrith/noverlap.

### Reversing to minimize yr count

So far, we have assumed that differences between related sequences are random and sequence-independent. This is not really true: in particular, vertebrates have a high rate of `cg` →`tg`/`ca` substitutions [36]. This suggests that words with fewer `yr`s may be better. As a first step in this direction, we reversed all words in our set if doing so would reduce the total `yr` count. Such reversal does not change the run-hitting probabilities calculated as above.

### Syncmer run-hitting probabilities

These were calculated using distance-distribution-parametrized-syncmer.py from https://tinyurl.com/frith-ISCB-V-Taiwan, which builds on “parametrized syncmer” concepts [9].

## Results

### Run-hitting probabilities

To start with our introductory example of `ry` versus `rr`, we calculated the probabilities of `ry` or `rr` occurring in runs of overlapping positions (fig. 1). `ry` has higher probability of occurring in a run of between 2 and at least 7 positions (fig. 2). This indicates that matches starting with `ry` between related sequences are more likely to occur than matches starting with `rr`.

**Figure 2:**
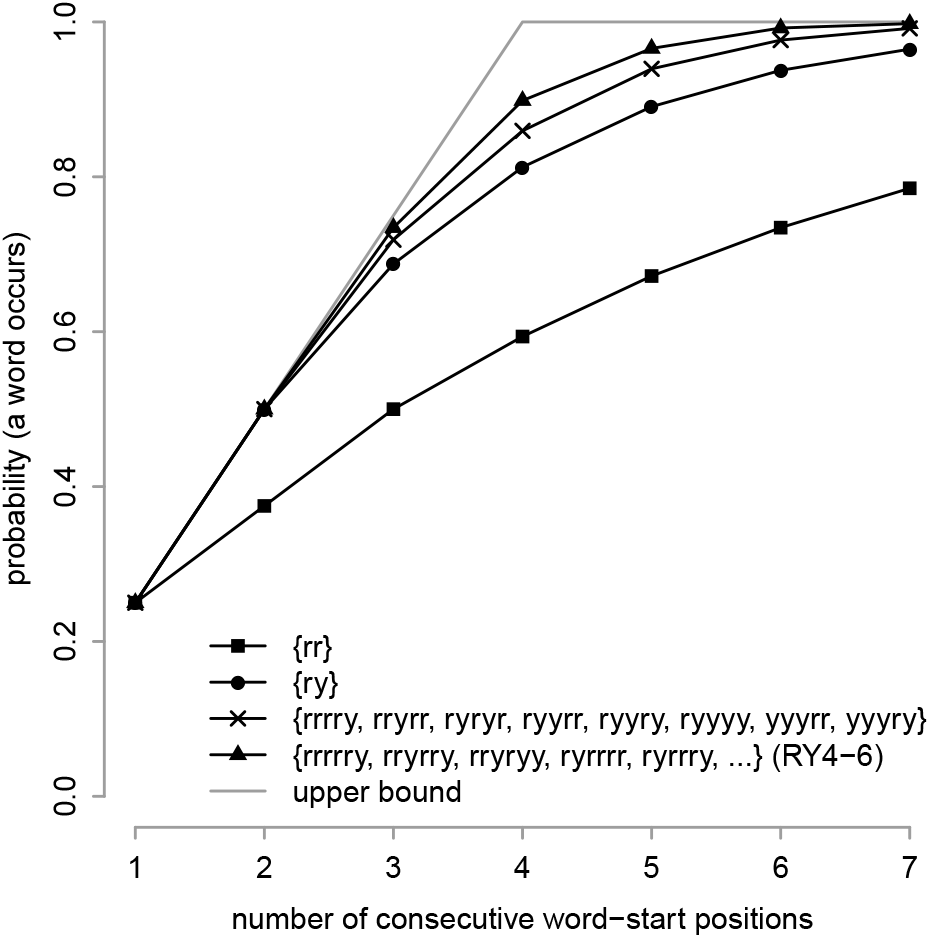
For each of four word sets, probability of occurring at least once in a sequence of random, independent, equally-probable `r` and `y`.

We then calculated run-hitting probabilities for some word sets found previously by heuristic criteria [12]. A set of eight length-5 words (fig. 2), which also samples 1 in 4 positions, has higher run-hitting probabilities than `ry`. Another set of sixteen length-6 words, named `RY4-6`, has even higher run-hitting ar probabilities (fig. 2). The eight 5-mers have the maximum average run-hitting probability, for run size 1 to *u*=7, among all sets of eight 5-mers (according to our exact optimization method). This shows that longer words enable higher run-hitting probabilities.

### Every *s*-th sampling is optimal

For comparing two sequences, one sampling method is to select every *s*-th position in one sequence, and every position in the other. This scheme samples 1 in *s* matches on average. It hits a run of *x < s* overlapping matches with probability *x/s*, and *x*≥ *s* matches with probability 1. Thus, it perfectly achieves the best-possible upper bound run-hitting probabilities.

This scheme has a disadvantage: it just samples one sequence. In constrast, if we use e.g. matches starting with a, we can rapidly skip all non-a positions in both sequences.

Another idea is to use matches whose start coordinates, in both sequences, are divisible by 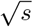, which samples 1 in *s* matches on average. This scheme hits runs of size 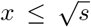 with optimal probability *x/s*. But for longer runs the hitting probability flatlines at 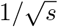.

### Optimizing run-hitting probabilities

We sought word sets whose run-hitting probabilities are as high as possible, for sparsity *s* = 4, 8, 16, and 32. If we use `r`/`y` words of length *k*, the number of words must be *n* = 2^*k*^*/s*. Longer words can achieve higher hitting probabilities, but are harder to optimize because there are more possible word sets. So we compromised by using heuristic optimization with *n* = 128 (table 1). We sought word sets with maximum average hitting probability, for run sizes from 1 to *u* = 2*s* − 1. This upper value *u* is arbitrary, but covers the range where the probabilities are most different from the upper bound (fig. 2, 3).^1^

**Table 1:** Sets of `r/y` words analyzed in this study

| name | sparsity | word<br>length | number<br>of words | min<br>separation | max | count of |  |
| --- | --- | --- | --- | --- | --- | --- | --- |
| | | | | | | $\mathbf{yr}$ | $\mathbf{ry}$ |
| *old* |  |  |  |  |  |  |  |
| RY4-6 | 4 | 6 | 16 | 2 | ∞ | 17 | 25 |
| RY8-8 | 8 | 8 | 32 | 3 | ∞ | 59 | 53 |
| RY16-8 | 16 | 8 | 16 | 6 | ∞ | 28 | 31 |
| RY32-10 | 32 | 10 | 32 | 9 | ∞ | 79 | 55 |
| *new* |  |  |  |  |  |  |  |
| RY4-9 | 4 | 9 | 128 | 2 | ∞ | 248 | 268 |
| RY4push | 4 | 9 | 128 | 3 | ∞ | 245 | 270 |
| RY4pull | 4 | 9 | 128 | 1 | 6 | 254 | 258 |
| RY8-10 | 8 | 10 | 128 | 4 | ∞ | 268 | 314 |
| RY16-11 | 16 | 11 | 128 | 7 | ∞ | 320 | 321 |
| RY32-12 | 32 | 12 | 128 | 10 | ∞ | 309 | 384 |

**Figure 3:**
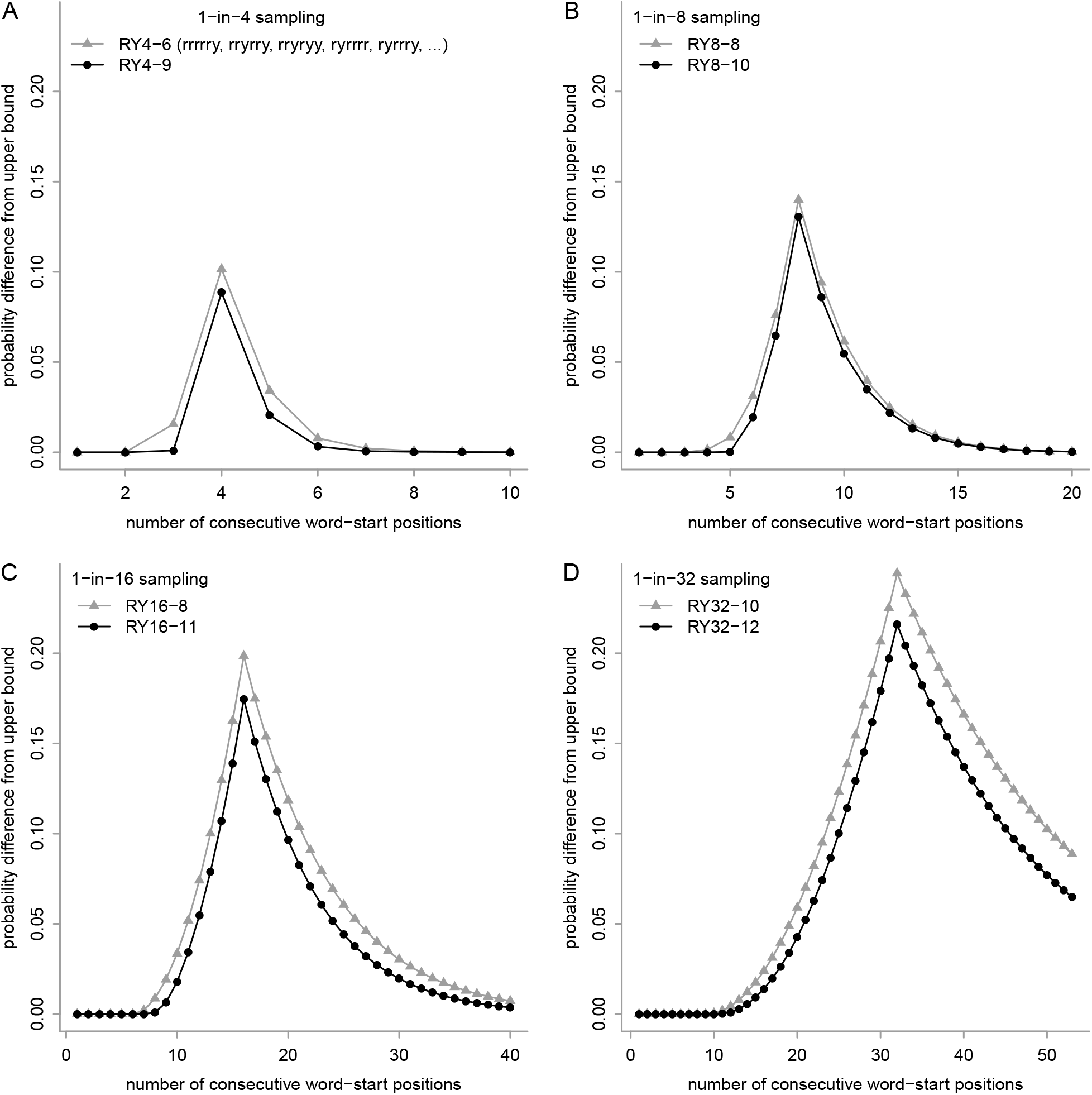
Run-hitting probabilities, for sampling schemes with sparsity: (**A**) 4, (**B**) 8, (**C**) 16, and (**D**) 32. For each run size (horizontal axis), the probability difference from the upper bound is shown (vertical axis). The gray triangles show the best schemes we found previously [12], and the black dots show new schemes. The triangles in panel **A** are equivalent to the triangles in fig. 2.

We found word sets with higher run-hitting probabilities, for each sparsity, than the best word sets we found previously (fig. 3). The names indicate sparsity and word length, e.g. `RY4-9` has sparsity 4 and word length 9. The improvement is small but clear.

### Optimizing a weighted average

By optimizing the average run-hitting probability, we attach equal importance to each run length, which is not necessarily correct. Actually, short runs are more important for comparing highly-diverged sequences, and long runs are more important for comparing highly-similar sequences. This is because runs of overlapping matches tend to be shorter in more-diverged sequences.

So we also tried optimizing a weighted average of the run-hitting probabilities, where the weight decreases by a factor *d* for each increment in run length. For sparsity 4, with *d* = 4 we found a set of length-9 words named `RY4push` (fig. 4). These words exactly achieve the upper bound run-hitting probability for run size ≤3. This means that these words cannot occur at more than 1 out of 3 consecutive positions, in other words, they sample positions with minimum separation 3. This is optimal for sparsity 4: a minimum separation of 4 would require that the spacing is always exactly 4, which is impossible in e.g. a homopolymer sequence like `rrrrrrrrrr`. A word set with guaranteed minimum separation has been termed a “polar set” [35].

**Figure 4:**
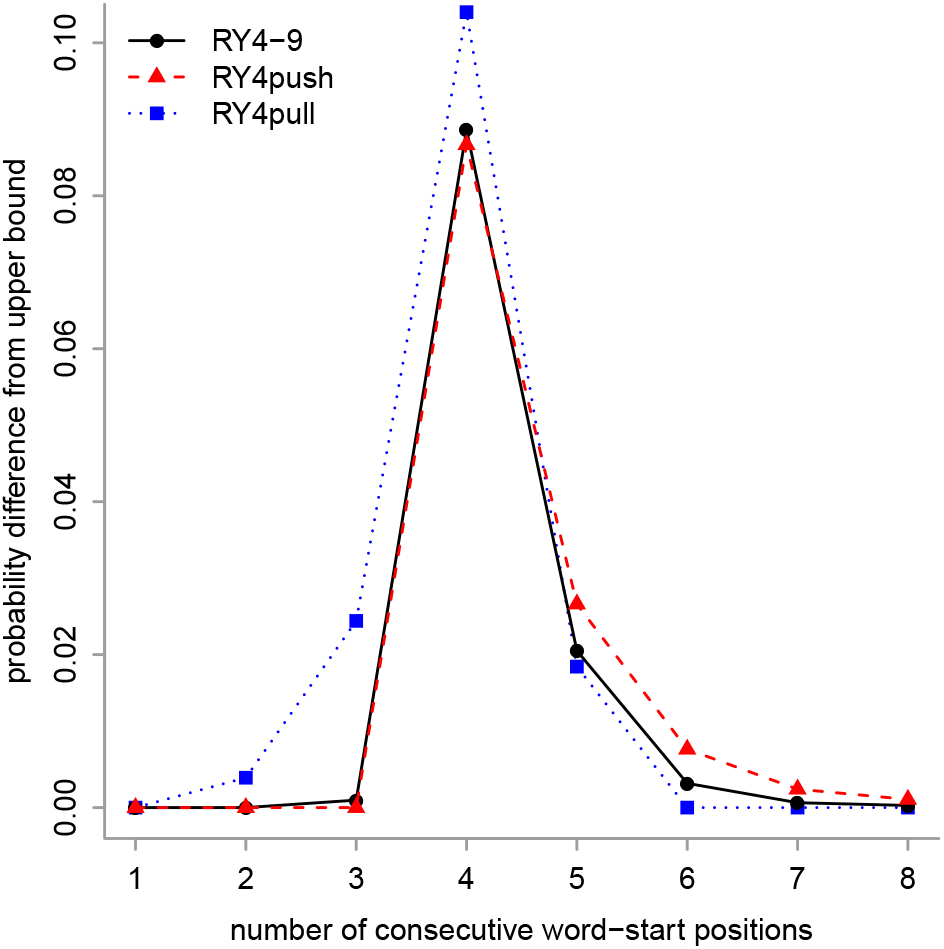
Run-hitting probabilities, for three 1-in-4 sampling schemes. For each run size (horizontal axis), the probability difference from the upper bound is shown (vertical axis). The black dots are the same as in fig. 3A.

We also tried attaching higher weight to longer runs. For sparsity 4, with *d* = ⅛ we found a set of length-9 words named `RY4pull` (fig. 4). These words achieve the upper bound run-hitting probability, which is 1, for run size ≥6. In other words, they sample positions with a maximum separation of 6. Since these words are guaranteed to occur in a finite sequence, they are an example of a universal hitting set [25].

In practice, `RY4-9` is likely near-optimal for both closely- and distantly-related sequences, because its run-hitting probabilities are never much lower than those of `RY4push` and `RY4pull` (fig. 4).

### One-hit specificity

Our approach increases the sensitivity of one-hit-based sequence comparison, but it’s important to consider specificity too. Sensitivity means probability of finding truly-related sequences, and specificity means probability of avoiding unrelated sequences. (The definition of “truly-related” depends on what we want to find, e.g. it could be homology or orthology.) For example, we can trivially increase sensitivity by using shorter matches, but that produces more matches between unrelated sequences, i.e. decreases specificity. Low seed specificity harms the run time of downstream seed-processing steps.

Is there any difference in specificity of `ry`- and `rr`- sampling? One way to measure specificity is by the number of sampled matches between unrelated random sequences: then there is no difference between `ry` and `rr`. However, one-hit seed-and-extend methods do not really attempt extension from every match: they avoid redundant extensions from overlapping matches. This suggests that two levels of specificity should be considered: all matches (which must be checked for overlap), and non-redundant matches (which trigger extension).

Let us investigate the number of sampled non-redundant matches between unrelated random sequences. The definition of non-redundant varies, but suppose that each match (of length *m*) is immediately extended into a maximally-long exact match, and subsequent matches in such extensions are skipped. Then, specificity can be measured by the number of sampled maximal exact matches between unrelated random sequences. The frequency of length-*y* maximal exact matches is proportional to *p*^*y*^ (where 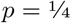 for DNA), and the probability of getting sampled is *H*_*y*−*m*+1_. So the expected fraction of maximal exact matches (of length ≥ *m*) that get sampled is

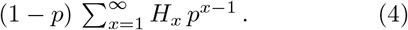

For example, `ry` samples a fraction 0.327 of maximal exact matches between unrelated random sequences, and `rr` samples 0.291. The upper bound, which applies to every 4-th sampling, is 0.332. So our approach has a downside: it maximizes non-redundant matches between unrelated sequences.

### Methods other than one-hit

Our sequence-sampling approach is beneficial not only for one-hit-based sequence comparison. Firstly, we conjecture that it maximizes the sensitivity of two-hit methods. Their sensitivity depends on finding two nearby, non-overlapping seeds: these seeds could be in the same run, or in separate runs (fig. 1). Our approach optimizes the probability of hitting separate runs, and of non-overlapping hits in one run (see the Supplement). Thus, it seems likely to improve the sensitivity of two-hit methods.

Our approach is also likely to increase the sensitivity of coverage-based sequence comparison. That is because it provably increases the expected number of letters covered by matches in a true alignment. This is proportional to the probability that one letter is covered, which was shown [30] to equal

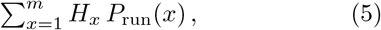

where *m* is the match length, and *P*_run_(*x*) is the probability that exactly *x* matches overlap one letter. Thus, higher run-hitting probabilities (*H*_*x*_) imply higher expected coverage, for any level of sequence divergence. Alignment-free methods are diverse, but a fundamental example is inferring the evolutionary distance between two sequences from the number of seed matches between them [20, 5]. To make this inference tight, we wish the variance in number of seed matches, for any given evolutionary distance, to be as low as possible [20]. Our sampling approach minimizes the variance of distance between sampled positions, which seems likely to reduce variance of match count.

### Comparison to syncmers

A recent sequence-sampling approach is syncmers [10]: how does it compare? Syncmers are a special case of word-based sampling. To define whether a length-*k* word is a syncmer, we first map each of its length-*j* subwords to an integer, by an arbitrary hash function. The number of subwords is *w* = *k*− *j* + 1. If the minimum *j*-mer is the first or last one in the *k*-mer, then the *k*-mer is a “closed syncmer”. If the minimum *j*-mer is the one at offset *t* (1 ≤*t* ≤*w*), then the *k*-mer is an “open syncmer with offset *t*”. Finally, a *d*-fold “down-sampled” open or closed syncmer is one where the hash function of the *k*-mer is ≤*h/d*, where *h* is the maximum possible hash.

To make syncmer properties simple, we shall assume the subword size *j* is large enough that tied hash values are rare. Then, open syncmers sample 1 in *w* positions, and closed syncmers sample 2 in *w* positions. Also, the run-hitting probabilities of open syncmers are maximized when *t* = [*w/*2] [30], so we shall assume this value for *t*.

The run-hitting probabilities of syncmers are shown in fig. 5. Closed syncmers are universal hitting sets: for sampling sparsity *s*, they guarantee to hit runs of size 2*s*≥ − 1. For shorter runs, however, their run-hitting probabilities are inferior. Open syncmers are polar word sets: they have minimum separation [*s/*2]. They are less sensitive than our word sets for sparsity 4 and 8, but more sensitive for sparsity 16 and 32. It is not too surprising that these open syncmers are more sensitive than our word sets, because they use longer words. The word length of open syncmers, for sparsity *s*, is *> s*. Thus, for sparsity 16 or 32, open syncmers can only be used with rather long matches, which have low sensitivity. Down-sampled open syncmers can use shorter words, but their run-hitting probabilities are inferior (fig. 5). Overall, open syncmers are a good heuristic for maximizing run-hitting probabilities.

**Figure 5:**
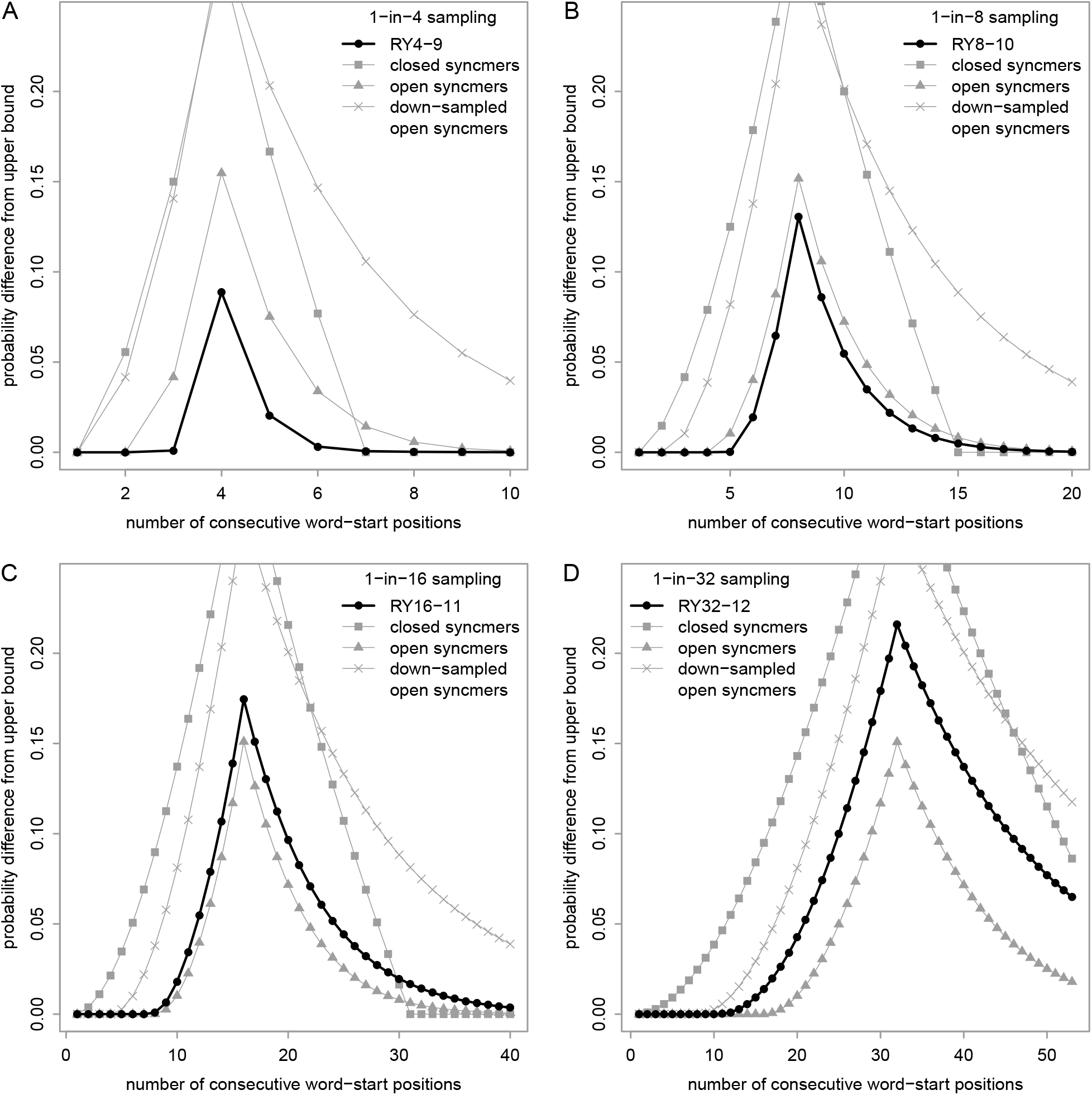
Run-hitting probabilities (difference from upper bound) for syncmers versus our word sets. The black dots are the same as in fig. 3. The down-sampled syncmers are 2-fold down-sampled.

### Sparsity in biological DNA

To see if our sampling methods are likely to work well for real DNA, we first checked the sparsity of our word sets in biological data. We checked human DNA (the first million non-`N` bases of hg38 chromosome 2), AT-rich *Plasmodium falciparum* DNA (the first million bases of NC 000521.4), and repetitive centromeric DNA (the 161 kb sequence 000736F from medaka strain Hd-rR 2.2.4 [13]). For the human and centromeric DNA, the words’ sparsity is close to the expected values of 4, 8, 16, and 32 (fig. 6). For the plasmodium DNA, some word sets are sparser than expected, but not wildly different. This suggests that our sampling schemes are applicable to a wide range of natural DNA.

**Figure 6:**
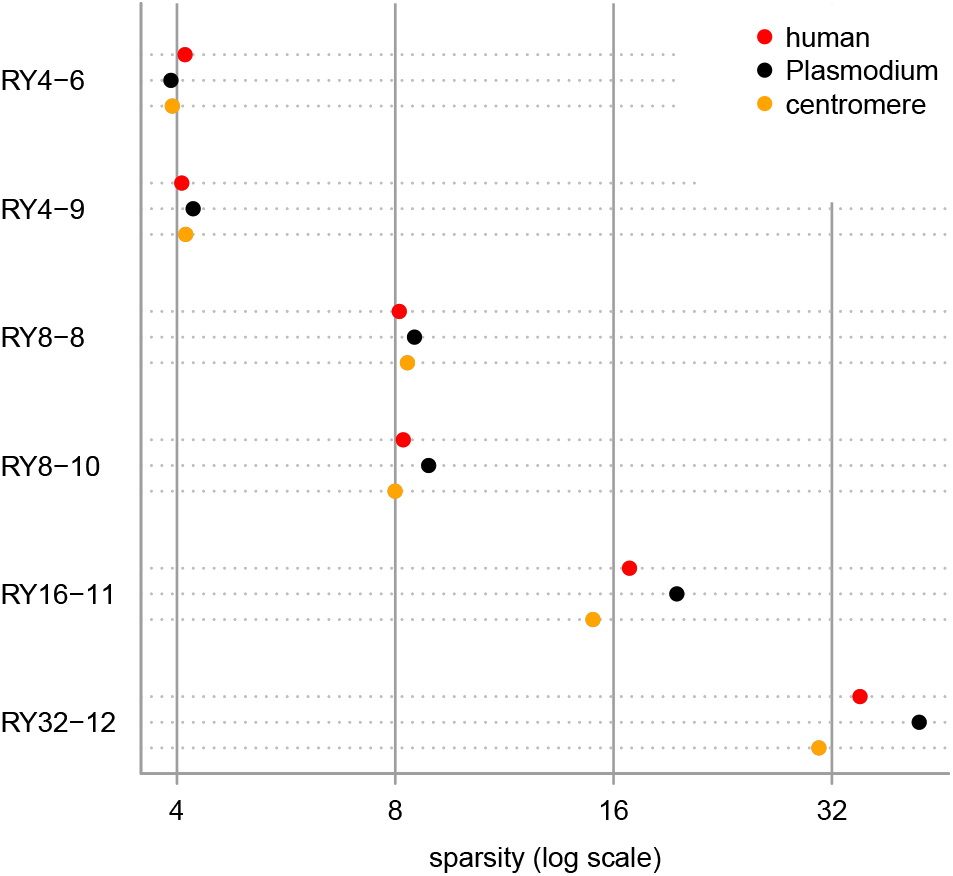
Sparsity of some sampling schemes in three biological DNA sequences.

### Specificity in biological DNA

We then measured specificity in real DNA, by counting the number of sampled matches between two unrelated sequences: 10^6^ bases of human DNA (the same as above) and 10^6^ bases of reversed (but not complemented) mouse DNA (the first million non-`N` bases of mm39 chromosome 1). Some sampling schemes find more matches than others (fig. 7), so they are less specific. For example, `RY4-6` finds more matches than `RY4-9`: this is because `RY4-6` includes the word `ryryry`, and that word alone gets most of the long matches found by `RY4-6`. This is presumably because `ryryry` samples simple repeats like `atatat`, which are frequent in real DNA. Similarly, `RY8-8` finds more matches than `RY8-10`, because `RY8-8` includes the repetitive words `yyyryyyr` and `yyryyryy`. By avoiding simple repeats, our new word sets are more specific than theoretically-optimal every *s*-th sampling.

**Figure 7:**
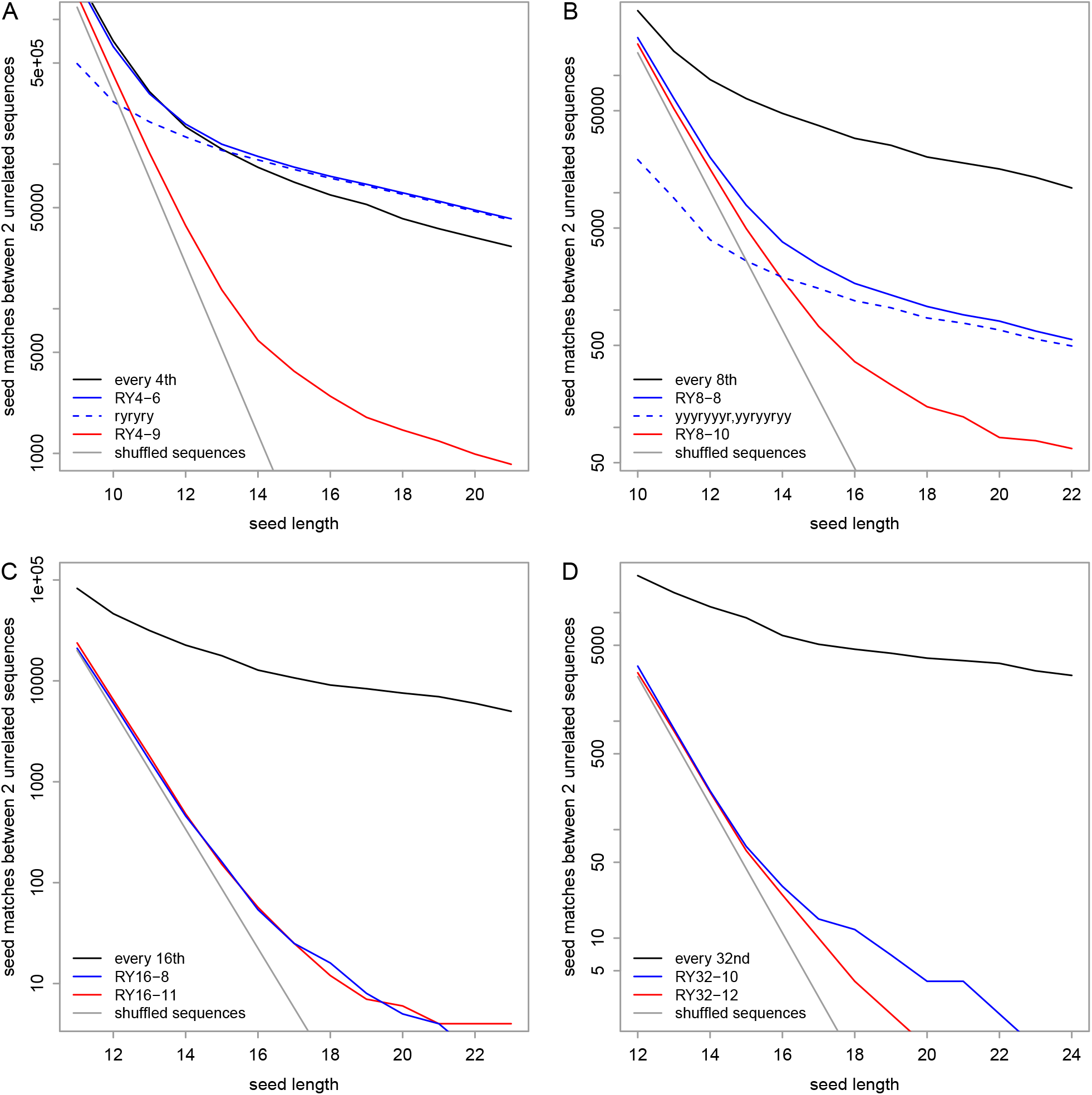
Specificity of some seed-sampling schemes, for mammal DNA. The vertical axis is the number of pair-wise seed matches between: 10^6^ bases of human DNA and 10^6^ bases of reversed (but not complemented) mouse DNA. In each panel, the gray line shows the expected number of hits if each sequence were shuffled.

### Sensitivity in biological DNA

Finally, we did one test of sensitivity in real DNA, by seeking short-range rearrangements in the gibbon genome relative to the human genome (e.g. fig. 8). Specifically, we sought rearrangements that may arise by short-range template switching during DNA replication, which are often overlooked [16].

**Figure 8:**
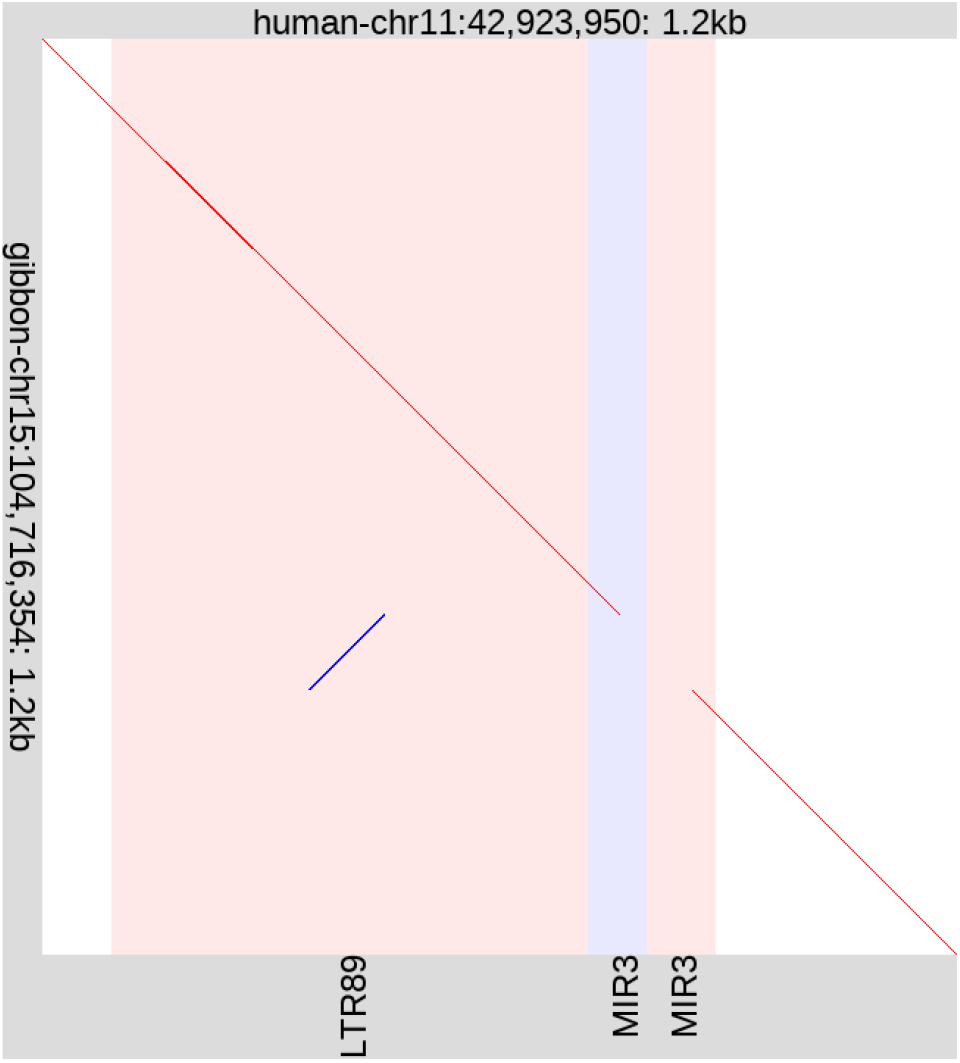
A short-range rearrangement in the gibbon genome relative to human. The diagonal lines show alignments: red = same strand, blue = opposite strand. The vertical stripes show repeat elements in the human sequence: red = forward-oriented, blue = reverse-oriented. By visually scanning this figure from top to bottom, we can see which part of the human sequence each part of the gibbon sequence comes from.

We used a two-step genome alignment method with the LAST software [11]. First, it finds and aligns similar segments of the two genomes. Second, it cuts these alignments down to a unique best alignment for each part of the gibbon genome. This is a reasonable way to find rearrangements that occurred in the gibbon lineage. The first step uses one-hit seed-and-extend, with variable-length seeds: at each sampled position in the gibbon sequence, it uses the shortest seed that occurs at most *m* times in human. Here we used exact-match seeds with *m* = 2.

As our ground truth, we used gibbon-human alignments found by the same method without any sampling. This will not be perfectly accurate, but it suffices to compare sampling methods’ abilities to find related segments. We defined “short range” rearrangements by arbitrary thresholds (see the Supplement), and deemed a sampling method to find a rearrangement if it finds one with the same start and end coordinates of the rearranged fragment.

In this test, the sensitivities of the sampling schemes agree with theoretical expectations. Firstly, every *s*-th sampling always has the highest sensitivity, e.g. every-16th sampling misses fewer rearrangements than `RY16-11` (fig. 9A). Secondly, our new word sets are always more sensitive than our old word sets (table 1), e.g. `RY4-9` misses fewer rearrangements than `RY4-6`. Furthermore, `RY4-6` is more sensitive than `RY`. Finally, the word sets are more sensitive when oriented to minimize `yr` count, in most cases.

**Figure 9:**
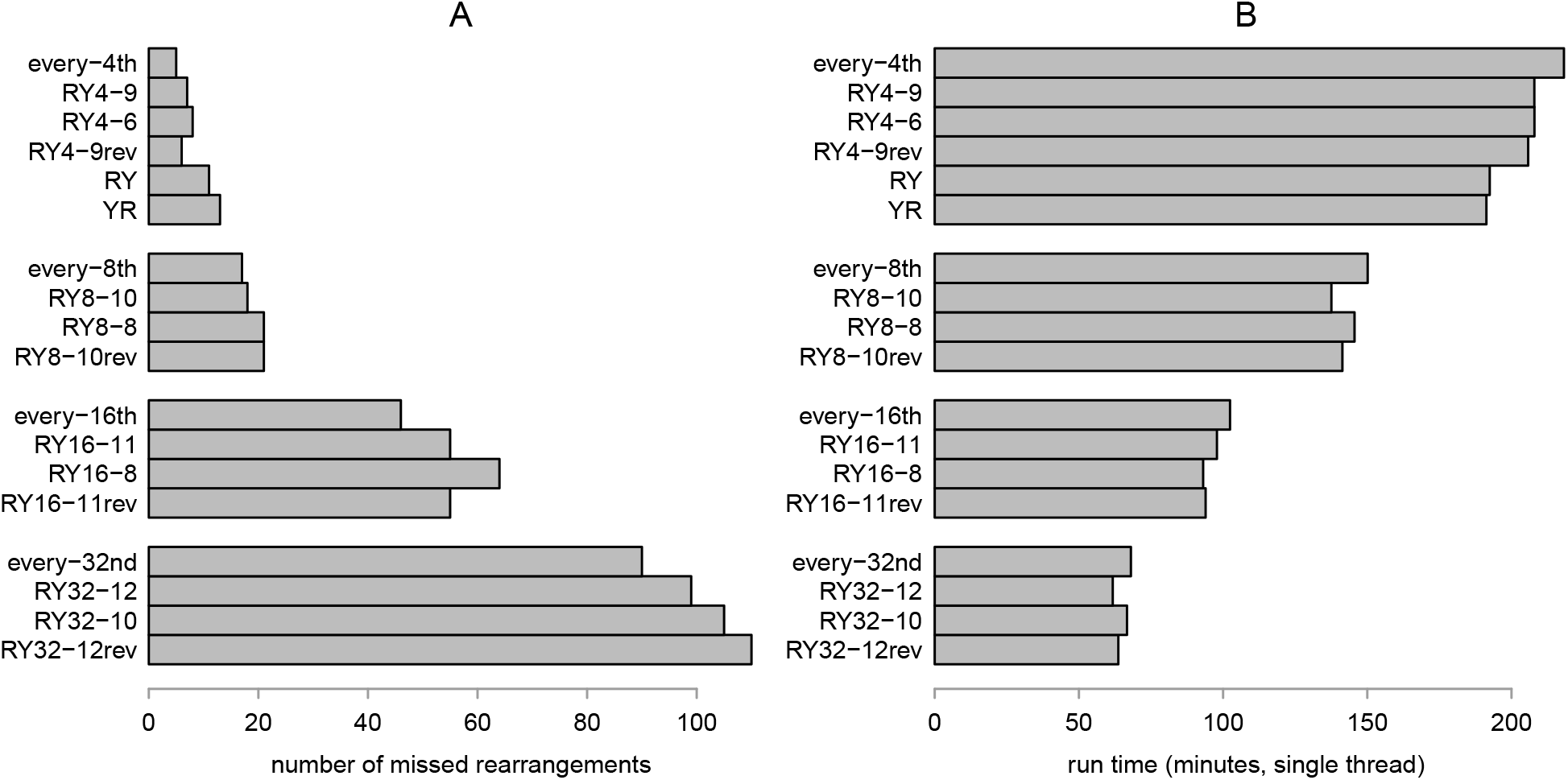
Sensitivity for finding short-range rearrangements, and run time, when aligning human and gibbon genomes using LAST with various seed-sampling schemes. The suffix rev indicates sets of reversed words, which thus maximize `yr` count.

The run times show the benefit of more-aggressive sampling (fig. 9B). Every *s*-th seeding used more memory, because it uses an index data-structure of all positions in the human genome, whereas word-based sampling just indexes the sampled positions. Among the 1-in-4 sampling schemes, RY and YR are fastest: we speculate this is because they have lower sensitivity so find fewer alignments.

## Discussion

We have presented an approach to sampling a sequence, which provably optimizes sensitivity for one-hit-based sequence comparison, for idealized random DNA. It is likely near-optimal for a wide range of alignment-based and alignment-free methods. For real biological DNA, our approach has close to the expected sampling rates, improves sensitivity in a limited test, and even improves specificity by avoiding simple repeats. Thus, our approach is promising for real DNA.

Our sampling method may have further applications. For example, various *k*-mer processing tasks are done by distributing *k*-mers into bins [24]. *k*-mers could be binned based on which (if any) of our words they contain. Because our words are evenly spaced, this binning is likely to be efficient and often put consecutive *k*-mers in the same bin.

Our sequence-sampling approach is a balance between universal hitting sets and polar sets. Universal hitting sets guarantee to sample a sufficiently-long sequence at least once, whereas polar sets have optimal hitting probabilities for sufficiently-short sequences which they sample at most once. It is impossible, however, for a word set to be both universal and polar. This is because a polar set with minimum separation *b* must exclude repeats (like `ryryry`) with period *< b*, but a universal set must include them. So it may be interesting to find “near-universal hitting sets” that hit every sequence except short-period repeats: such a set could perhaps be polar, with the specificity benefit of avoiding simple repeats.

There are several other directions for future advances. Firstly, it would be useful to find better-optimized word sets than we found by simulated annealing, with longer words. Related to this, it would be interesting to know if optimal or near-optimal word sets have any simple properties. It would be useful to optimize sampling of inexact seeds [12], which we feel are under-used, though they benefit alignment-based and alignment-free methods (e.g. [17, 20, 26]). In particular, transition seeds, which tolerate frequently-occurring `a` ↔ `g` and `c` ↔ `t` substitutions, are greatly neglected and promising for improved sequence comparison [22]. It would also be useful to optimize protein sequence sampling, perhaps using a reduced alphabet. Finally, it may be useful to optimize sampling of specific sequences, such as a reference human genome [7, 35]: it will be interesting to see how much this improves over sampling methods that are optimal for random sequences.

We have clarified how to do optimal sampling of a sequence, which is a fundamental way to rapidly analyze big sequence data. At the same time, we have shown that random sequences of independent letters are surprisingly interesting.

## Acknowledgements

We thank Anish Shrestha, Kiyoshi Asai, and Johan Nyström-Persson for helpful comments, and Animesh Awasthi for a comparison of sequence-sampling methods which motivated this work.

## Footnotes

1 Actually, we set a hard upper limit on *u*: *u* + *k* − 1 ≤64, because our software does not handle higher values.

