## Supplement for "How to optimally sample a sequence for rapid analysis"

December 4, 2022

### Run-hitting probabilities & distances

For a sequence-sampling scheme, we can calculate the probability $H_x$ of hitting a size-$x$ run, from the probabilities $D_i$ of distance $i$ between consecutive sampled positions. Consider a size-$x$ run at a random location in the middle of a long sequence. Get the nearest sampled position before the run start, and the nearest sampled position at or after the run start. The distance between these two sampled postions equals some integer $i \geq 1$ with probability proportional to $iD_i$. So this probability is $iD_i/s$, where $s$ is the sparsity or expected distance $\sum_{i=1}^{\infty} iD_i$. The probability that the run is hit, given $i$, is $\min(x/i, 1)$. Therefore

$$H_x = \sum_{i=1}^{\infty} \frac{iD_i}{s} \min(x/i, 1) \tag{S1}$$

$$= \left(\sum_{i=1}^{x} iD_i + \sum_{i=x+1}^{\infty} xD_i\right)/s \tag{S2}$$

$$= \left(x - \sum_{i=1}^{x} (x-i)D_i\right)/s\,. \tag{S3}$$

### Inverse formula

We can also calculate $D_i$ from $H_x$:

$$D_x = (2H_x - H_{x-1} - H_{x+1})s\,, \tag{S4}$$

where $H_0 = 0$. To verify Eq (S4), substitute the right-hand side of Eq (S3) into the right-hand side of Eq (S4), and observe that it equals $D_x$.

### Distance & average hitting probability

Here, we show that maximizing the average run-hitting probability is equivalent to minimizing the variance of distance between consecutive sampled positions.

First, define $L_x$ to be the probability of sampling the final position, and not any earlier position, of a size-$x$ run. It follows that

$$H_x = \sum_{i=1}^{x} L_i \tag{S5}$$

and

$$L_x = H_x - H_{x-1}\,. \tag{S6}$$

By combining this with Eq (S4), we get

$$D_x = (L_x - L_{x+1})s\,. \tag{S7}$$

It's also useful to define $\overline{H}_x = 1 - H_x$, so that

$$L_x = \overline{H}_{x-1} - \overline{H}_x\,. \tag{S8}$$

Next, as a stepping stone to the variance,

$$\mathrm{Var}[D] = \left(\sum_{i=1}^{\infty} i^2 D_i\right) - s^2\,, \tag{S9}$$

we first calculate

$$\sum_{i=1}^{u} i^2 D_i/s$$

$$= \sum_{i=1}^{u} i^2(L_i - L_{i+1}) \tag{S10}$$

$$= \sum_{i=1}^{u} (2i-1)L_i - u^2 L_{u+1} \tag{S11}$$

$$= \sum_{i=1}^{u} (2i-1)(\overline{H}_{i-1} - \overline{H}_i) - u^2 L_{u+1} \tag{S12}$$

$$= 2\sum_{i=1}^{u} \overline{H}_i - (2u+1)\overline{H}_u + 1 - u^2 L_{u+1} \tag{S13}$$

$$= 2\sum_{i=1}^{u} \overline{H}_i - (u+1)^2\overline{H}_u + 1 + u^2\overline{H}_{u+1}\,. \tag{S14}$$

Steps (S11) and (S13) are examples of summation by parts (`https://en.wikipedia.org/wiki/Summation_by_parts`).

Now we show that $\overline{H}_x$ is exponentially decreasing in $x$. This implies that all terms in (S14) converge. The argument is as follows: given $x$ consecutive positions, for a hit to occur we require that a $k$-length word in the $x+k-1$ sequence positions is sampled. We can find $\gamma(x) = \lfloor \frac{x+k-1}{k} \rfloor$ non-overlapping $k$-length words. Since these are non-overlapping, each such word is independently sampled with probability $\frac{1}{s}$. Thus the probability that one of these non-overlapping words is sampled is $1-(1-\frac{1}{s})^{\gamma(x)}$ by independence. This is upper bounded by $H_x$, which considers all words and not just these particular non-overlapping words. This gives us the exponentially decreasing upper bound on $\overline{H}_x$,

$$\overline{H}_x \leq \left(1 - \frac{1}{s}\right)^{\gamma(x)} . \qquad \text{(S15)}$$

This argument works for all word-based sampling schemes, and can even be extended to sequence sampling schemes such as minimizers after a small modification. We can now see that

$$\lim_{u\to\infty} \left(\sum_{i=1}^{u} i^2 D_i/s - 2\sum_{i=1}^{u} \overline{H}_i - 1\right) = -(u+1)^2\overline{H}_u + u^2\overline{H}_{u+1} \to 0 , \qquad \text{(S16)}$$

where we used the above fact that $\overline{H}_u \to 0$ exponentially fast. The limit shows that

$$\mathrm{Var}[D] = \left(2\sum_{i=1}^{\infty} \overline{H}_i + 1 - s\right)s . \qquad \text{(S17)}$$

This means that minimizing $\mathrm{Var}[D]$, for fixed $s$, is equivalent to minimizing the sum of $\overline{H}_i$. This is approximately equivalent to maximizing a finite sum $\sum_{i=1}^{u} H_i$ (or average) of $H_i$.

### Two-hit probabilities

Our sequence-sampling approach seems likely to improve the sensitivity of two-hit sequence comparison methods. In particular, it optimizes the probability of sampling two non-overlapping length-$m$ seeds within one run of $x$ unmutated seeds. This probability is:

$$H_{2,x} = \sum_{i=1}^{x-m} L_i H_{x-m-i+1} \qquad \text{(S18)}$$

$$= \sum_{i=1}^{x-m} (H_i - H_{i-1}) H_{x-m-i+1} . \qquad \text{(S19)}$$

The formula follows by conditioning on the position $i$ of the first seed, which introduces an $L_i$ term. Subject to this condition, there are $x-m-i+1$ possible consecutive positions for seeds that do not overlap the first seed. The characters in these positions are independent of the seed match at $i$, so we just get $H_{x-m-i+1}$, the probability of seed occurring in the remaining consecutive positions.

If we increase the $H_i$ values, $H_{2,x}$ never decreases. This is because the partial derivative of $H_{2,x}$ with repect to $H_i$ is $2H_{x-m-i+1} - 2H_{x-m-i}$, which is $\geq 0$. Thus, maximizing $H_i$ implies maximizing $H_{2,x}$.

### Simulated annealing

To find a set of words with high run-hitting probabilities, we used a simulated annealing method. Starting with a random set of $n$ words, we iterate the following procedure. We randomly pick one word in the set and one word not in the set, and try exchanging them. If this increases the objective function (average run-hitting probability), we accept the exchange. If it decreases the objective function by an amount $z$, then we randomly accept or reject, with acceptance probability $e^{-z/t}$, where $t$ is a parameter called "temperature". Thus, low temperature rarely decreases the objective function, while high temperature can escape local optima. We start by setting $t$ equal to: the objective function's upper bound minus the objective function's intial value. We then multiply $t$ by something like 0.99999999 at each iteration, until the words "freeze".

### Gibbon-human genome alignment

The gibbon (nomLeu3) and human (hg38.analysisSet) genomes were aligned with LAST version 1406. First, we found the rates of deletion, insertion, and each kind of substitution between the genomes:

```
lastdb -uRY16 myDB human.fa

last-train --revsym myDB gibbon.fa > train.out
```

`lastdb` makes an index data-structure named `myDB` for the human genome, and `last-train` infers the rates. The `-uRY16` option, in this version of LAST, specifies `RY16-8` sampling. The sampling scheme has very little effect on the result. The resulting substitution score matrix is

```
     A    C    G    T
A    5  -15   -9  -16
C  -15    7  -13  -10
G  -10  -13    7  -15
T  -16   -9  -15    5
```

where rows correspond to human bases and columns to gibbon bases. The cost of a length-$g$ insertion or deletion is $34 + g$. We then aligned the genomes with word-based sampling like this:

```
lastdb -dwords myDB human.fa

lastal -C2 -m2 --split -p train.out myDB
                                  gibbon.fa > out.maf
```

Here, *words* was replaced by a word list such as `RRRY,RYRR,RYYR,YYYR`. We aligned the genomes with every $s$-th sampling like this:

```
lastdb -m1 denseDB human.fa

lastal -C2 -m2 -ks --split -p train.out denseDB
                                  gibbon.fa > out.maf
```

Here, $s$ was replaced by 4, 8, 16, or 32, or by 1 for no sampling.

### Short-range rearrangements

We used the following arbitrary thresholds to define "short-range" switch-like rearrangements. From the whole-genome alignment, we have a set of alignments between gibbon and human segments, with no overlap in gibbon. In each gibbon chromosome, a group of three consecutive alignments $(X, Y, Z)$ was deemed a switch-like rearrangement if:

- They align to the same human chromosome.
- $X$ and $Z$ align to the same human strand, and $Y$ to the opposite strand.
- There are at most 10 unaligned gibbon base-pairs between $X$ and $Y$, and between $Y$ and $Z$.
- There are at most 1000 human base-pairs between Y and the end of $X$, or between $Y$ and the start of $Z$.
- There are at most 1000 human base-pairs between the end of $X$ and the start of $Z$.
- In human, the number of base-pairs spanned by $Y$ is at most double the number of base-pairs between end of $X$ and the start of $Z$.

In our ground-truth rearrangements, the median number of human base-pairs spanned by $Y$ was 127.

### Word sets

RY4-9

rrrrrrrry rrrrrryry rrrrryrrr rrrrryrry
rrrrryyrr rrrrryyry rrrrryyyr rrrrryyyy
rrryryrrr rrryryrry rrryryyrr rrryryyry
rrryryyyr rrryryyyy rryrrrrrr rryrrrrry
rryrrryry rryrryrrr rryrryrry rryrryryr
rryrryyrr rryrryyry rryrryyyy rryryryry
rryyryrrr rryyryrry rryyryyry rryyryyyy
rryyyyrrr rryyyyrry rryyyyryr rryyyyyrr
ryrrrryrr ryrrrryry ryrrrryyr ryrrrryyy
ryrrryrrr ryrrryrry ryrrryyrr ryrrryyry
ryrrryyyr ryrrryyyy ryrryryyr ryrryryyy
ryrryyyrr ryrryyyry ryryryrrr ryryryrry
ryryryyrr ryryryyry ryryryyyr ryryryyyy
ryyrrrrrr ryyrrrrry ryyrrrryr ryyrrryrr
ryyrrryry ryyrrryyr ryyrrryyy ryyrryrrr
ryyrryrry ryyrryryr ryyrryyrr ryyrryyry
ryyrryyyr ryyrryyyy ryyryryrr ryyryryry
ryyryryyr ryyryryyy ryyryyrrr ryyryyrry
ryyryyyrr ryyryyyry ryyyryrrr ryyyryrry
ryyyryyrr ryyyryyry ryyyryyyr ryyyryyyy
ryyyyryyr ryyyyryyy ryyyyyryr ryyyyyyrr
ryyyyyyyy yryrrrrrr yryrrrrry yryrrryry
yryrryrrr yryrryrry yryrryryr yryrryyrr
yryrryyry yryrryyyy yryyryrrr yryyryrry
yryyryyry yryyryyyy yryyyyrrr yryyyyrry
yryyyyryr yryyyyyrr yyrrrryyr yyrrrryyy
yyrryryyr yyrryryyy yyyrrrrrr yyyrrrrry
yyyrrrryr yyyrrryrr yyyrrryry yyyrrryyr
yyyrrryyy yyyrryrrr yyyrryrry yyyrryryr
yyyrryyrr yyyrryyry yyyrryyyr yyyrryyyy
yyyryryrr yyyryryry yyyryryyr yyyryryyy
yyyyyryyr yyyyyryyy yyyyyyryr yyyyyyyrr

RY8-10

rrrryrrrry rrrryrrryr rrrryrrryy rrrryryrry
rrrryryryy rrrryyrryy rrrryyyryy rrryrrrrrr
rrryrrrrry rrryrryrry rrryrryryy rrryyrrrry
rrryyrrryr rrryyrrryy rrryyrryry rrryyryrry
rrryyryryy rrryyyrrrr rrryyyrrry rrryyyrryy
rrryyyyryy rryryrrrrr rryryrrrry rryryrrryr
rryryrrryy rryryryrrr rryryryrry rryryryryr
rryryryryy rryryyrryy rryryyyryy rryyrrrrry
rryyrryrrr rryyrryrry rryyryrrrr rryyryrrry
rryyryryrr rryyryryry rryyryyryy rryyyrryrr
rryyyrryry rryyyryrry rryyyryryy rryyyyrrrr
rryyyyrrry rryyyyrryr rryyyyrryy rryyyyryrr
rryyyyryry rryyyyyrry rryyyyyryy rryyyyyyry
ryrryrrrrr ryrryrrrry ryrryrrryr ryrryrrryy
ryrryryrry ryrryryryy ryrryyrryy ryrryyyryy
ryryyrrrry ryryyrrryr ryryyrrryy ryryyrryry
ryryyryrry ryryyryryy ryryyyrryy ryryyyyryy
ryyrrrrrrr ryyrrrrrry ryyryyrryy ryyryyyryy
ryyyryrrrr ryyyryrrry ryyyryryrr ryyyryryry
ryyyyyrrrr ryyyyyrrry ryyyyyryrr ryyyyyryry
ryyyyyyrrr ryyyyyyrry ryyyyyyyry yrryyrrrry
yrryyrrryr yrryyrrryy yrryyrryry yrryyryrry
yrryyryryy yrryyyrrrr yrryyyrrry yrryyyrryy
yrryyyyryy yryyrrrrry yryyrryrrr yryyrryrry
yryyryrrrr yryyryrrry yryyryryrr yryyryryry
yryyyrrrrr yryyyrrrry yryyyrrryr yryyyrrryy
yryyyrryrr yryyyrryry yryyyryrry yryyyryryy
yryyyyrrrr yryyyyrrry yryyyyrryr yryyyyrryy
yryyyyryrr yryyyyryry yryyyyyrry yryyyyyryy
yryyyyyyry yyryyrrrry yyryyrrryr yyryyrrryy
yyryyrryry yyryyryrry yyryyryryy yyryyyrryy
yyryyyyryy yyyyyyyrrr yyyyyyyrry yyyyyyyyry

RY16-11

rryryrrrrrr rryryrrrryr rryryrrryrr rryryrryrrr
rryryrryrry rryryryrrrr rryryryrryr rryryryryrr
rryryryyrrr rryryryyrry rryryyrrrrr rryryyrrryr
rryryyrryrr rryryyryrry rryryyyrrrr rryryyyrryr
rryryyyryrr rryryyyyrrr rryryyyyrry rryyrrrrrrr
rryyrrrrrry rryyrrrrryr rryyrrrryrr rryyrrryrrr
rryyrrryrry rryyrryrrrr rryyrryrrry rryyrryrryr
rryyryrrrrr rryyryrrrry rryyryrrryr rryyryrryrr
rryyryryrrr rryyryryrry rryyryryyrr rryyryyrrrr
rryyryyrrry rryyryyrryr rryyryyyrrr rryyryyyrry
rryyryyyyrr rryyyrrrrrr rryyyrrrrry rryyyrrrryr
rryyyrrryrr rryyyrryrrr rryyyrryrry rryyyryrrrr
rryyyryrrry rryyyryrryr rryyyryyrrr rryyyryyrry
rryyyryyyrr rryyyyrrrrr rryyyyrrrry rryyyyrrryr
rryyyyrryrr rryyyyryrrr rryyyyryrry rryyyyryyrr
rryyyyyrrrr rryyyyyrrry rryyyyyrryr rryyyyyryrr
rryyyyyyrry ryryryryyrr ryryryyyrrr ryryryyyrry
ryryryyyyrr ryryyryrrrr ryryyryrrry ryryyryryrr
ryryyryyrrr ryryyryyrry ryryyryyyrr ryryyyryyrr
ryryyyyryrr ryryyyyyrrr ryryyyyyrry ryyryyryrrr
ryyryyryrry ryyryyyryrr ryyyryryrrr ryyyryryrry
ryyyryryyrr ryyyryyryrr ryyyyryryrr ryyyyyryyrr
ryyyyyyrrrr ryyyyyyrrry ryyyyyyryrr ryyyyyyyrrr
ryyyyyyyrry ryyyyyyyyrr yryryryryrr yryryyryrry
yryryyyryrr yryyryryyrr yryyryyyyrr yryyyryyyrr
yryyyyryyrr yryyyyyryrr yryyyyyyrry yyryryryrrr
yyryryryrry yyryryryyrr yyryryyyrrr yyryryyyrry
yyryryyyyrr yyryyryryrr yyryyryyrrr yyryyryyrry
yyryyryyyrr yyryyyryyrr yyryyyyryrr yyryyyyyrrr
yyryyyyyrry yyyryyyryrr yyyryyyyrrr yyyryyyyrry
yyyyryryyrr yyyyryyryrr yyyyryyyrrr yyyyryyyrry
yyyyyryryrr yyyyyyryyrr yyyyyyyryrr yyyyyyyyyrr

RY32-12

rrrrrrrryyyr rrrrrrrryyyy rrrrrryrryyy rrrrrryryyyr
rrrrryrrryyy rrrrryryyyyr rrrrryryyyyy rrrrryyryyyr
rrrrryyryyyy rrrryryrryyy rrrryryryyyr rrrryryryyyy
rrrryryyryyy rrrryyrryyyr rrrryyrryyyy rrrryyryryyy
rrryrrrryyyr rrryrrrryyyy rrryrryryyyr rrryrryryyyy
rrryyrrrryyy rrryyrrryyyr rrryyryrryyy rrryyryyryyy
rryrrrrrryyy rryrrrrryyyr rryrrryryyyr rryrrryryyyy
rryrrryyryyy rryrryrrryyy rryrryrryyyr rryrryrryyyy
rryrryyrryyy rryrryyryyyr rryrryyryyyy rryryrrrryyy
rryryrrryyyr rryryrrryyyy rryryrryryyy rryryryrryyy
rryryryryyyr rryryryryyyy rryryyrrryyy rryryyrryyyr
rryryyrryyyy rryyrrrrryyy rryyrrryryyy rryyrrryyyyr
rryyrrryyyyy rryyrryryyyr rryyrryryyyy rryyryryryyy
rryyryyrryyy ryrrrrrrryyy ryrrrrryyyyr ryrrrrryyyyy
ryrrrryrryyy ryrrrryryyyr ryrrrryryyyy ryrrrryyryyy
ryrrryrrryyy ryrrryrryyyr ryrrryrryyyy ryrrryryryyy
ryrrryyrryyy ryrryrrrryyy ryrryrryryyy ryrryryrryyy
ryrryryryyyr ryrryryryyyy ryrryryyryyy ryrryyryryyy
ryryrrryryyy ryryrryrryyy ryryrryyryyy ryryryrrryyy
ryryryryryyy ryryryyrryyy ryryryyryyyr ryryryyryyyy
ryryyrrrryyy ryryyrryryyy ryryyryrryyy ryryyryryyyr
ryryyryryyyy ryryyryyryyy ryyrrrrrryyy ryyrrrryryyy
ryyrrryrryyy ryyrrryyryyy ryyrryrrryyy ryyrryrryyyr
ryyrryrryyyy ryyrryryryyy ryyrryyrryyy ryyrryyryyyr
ryyrryyryyyy ryyryrrrryyy ryyryrrryyyr ryyryrrryyyy
ryyryrryryyy ryyryryrryyy ryyryryyryyy ryyryyrrryyy
ryyryyryryyy yrrrrryrryyy yrrrrryryyyr yrrrryrrryyy
yrrryryrryyy yrrryryyryyy yrrryyryryyy yrryyrrrryyy
yrryyrrryyyr yrryyryrryyy yrryyryyryyy yryrrrrrryyy
yryrrrrryyyr yryrrryyryyy yryrryrrryyy yryrryyrryyy
yryryrrrryyy yryryrryryyy yryryryrryyy yryryyrrryyy
yryyrrrrryyy yryyrrryryyy yryyryryryyy yryyryyrryyy

```
RY4push
rrrrryrrr rrrrryrry rrrrryryr rrrrryryy
rrrrryyrr rrrrryyry rrrrryyyy rrrryyyrr
rrrryyyry rryrrrrry rryrryrrr rryrryrry
rryrryryr rryryrrrr rryryrrry rryryrryr
rryryrryy rryryryry rryryyrrr rryryyrry
rryryyryr rryryyryy rryryyyrr rryryyyry
rryryyyyr rryryyyyy rryyyyrrr rryyyyrry
rryyyyryr rryyyyryy ryrrrrrrr ryrrrrrry
ryrrrryrr ryrrrryry ryrrrryyr ryrrryrrr
ryrrryrry ryrrryryr ryrrryryy ryrrryyrr
ryrrryyry ryrrryyyy ryrryyrrr ryrryyrry
ryrryyryr ryrryyryy ryrryyyrr ryrryyyry
ryrryyyyy ryryryrrr ryryryrry ryryryyrr
ryryryyry ryryryyyy ryyrrrrrr ryyrrrrry
ryyrrryry ryyrryrrr ryyrryrry ryyrryryr
ryyrryryy ryyrryyrr ryyrryyry ryyrryyyy
ryyryrrrr ryyryrrry ryyryrryy ryyryryry
ryyryyrrr ryyryyrry ryyryyryr ryyryyryy
ryyryyyrr ryyryyyry ryyryyyyy ryyyyyrrr
ryyyyyrry ryyyyyryr ryyyyyryy ryyyyyyrr
ryyyyyyry yrrrryyyy yrrryyyrr yrrryyyry
yryryyyrr yryryyyry yyrrrryrr yyrrrryry
yyrrrryyr yyrrryrrr yyrrryrry yyrrryyrr
yyrrryyry yyrrryyyy yyrryyyrr yyrryyyry
yyryrryrr yyryrryry yyryryrrr yyryryrry
yyryryyrr yyryryyry yyryryyyy yyryyyyrr
yyryyyyry yyyrrrrrr yyyrrrrry yyyrrryry
yyyrryrrr yyyrryrry yyyrryryr yyyrryryy
yyyrryyrr yyyrryyry yyyrryyyy yyyryrrrr
yyyryrrry yyyryrryy yyyryryry yyyryyrrr
yyyryyrry yyyryyryr yyyryyryy yyyryyyrr
yyyryyyry yyyryyyyy yyyyyyyrr yyyyyyyry

RY4pull
rrrrrrrrr rrrrrryrr rrrrrryry rrrrrryyr
rrrrrryyy rrryryrrr rrryryyrr rrryryyry
rrryryyyr rrryryyyy rryrrrrrr rryrrryrr
rryrrryry rryrrryyr rryrryrrr rryrryrry
rryrryryr rryrryyrr rryrryyry rryrryyyr
rryrryyyy rryryryyr rryryryyy rryyryrrr
rryyryyrr rryyryyry rryyryyyr rryyryyyy
rryyyryrr rryyyyrrr rryyyyyrr rryyyyyyr
ryrrrryrr ryrrrryry ryrrrryyr ryrrrryyy
ryrrryyyr ryrrryyyy ryrryryyr ryrryryyy
ryryrryrr ryryrryry ryryrryyr ryryrryyy
ryryryrrr ryryryrry ryryryryr ryyrrrrrr
ryyrrrrry ryyrrryrr ryyrrryry ryyrrryyr
ryyrrryyy ryyrryrrr ryyrryrry ryyrryryr
ryyrryyrr ryyrryyry ryyrryyyr ryyrryyyy
ryyryryrr ryyryryry ryyryryyr ryyryryyy
ryyyrrrrr ryyyrrrry ryyyrryrr ryyyrryry
ryyyrryyr ryyyrryyy ryyyryryr ryyyryyrr
ryyyryyry ryyyryyyr ryyyryyyy ryyyyryrr
ryyyyryry ryyyyryyr ryyyyryyy ryyyyyyyr
yrrrrryrr yrrrrryry yrrrrryyr yrrrrryyy
yrryryrrr yryrrrrrr yryrrryrr yryrrryry
yryrrryyr yryryryyr yryryryyy yryyryrrr
yryyryyrr yryyryyry yryyryyyr yryyryyyy
yryyyryrr yryyyyrrr yryyyyyrr yryyyyyyr
yyrrrryrr yyrrrryry yyrrrryyr yyrrrryyy
yyrryryyr yyrryryyy yyryrryrr yyryrryry
yyryrryyr yyryrryyy yyryryrrr yyyrrryrr
yyyrrryry yyyrrryyr yyyrrryyy yyyryryyr
yyyryryyy yyyyrrrrr yyyyrryrr yyyyrryry
yyyyrryyr yyyyrryyy yyyyyryrr yyyyyryry
yyyyyryyr yyyyyryyy yyyyyyyyr yyyyyyyyy
```
